# Non-CG methylation is superior to CG methylation in genome regulation

**DOI:** 10.1101/2020.03.04.971267

**Authors:** Katherine Domb, Aviva Katz, Rafael Yaari, Efrat Kaisler, Vu Hoang Nguyen, Uyen Vu Thuy Hong, Ofir Griess, Karina Gitin Heskiau, Nir Ohad, Assaf Zemach

**Affiliations:** School of Plant Sciences and Food Security, Tel-Aviv University, Tel- Aviv 69978, Israel

**Author notes:** Equal contribution (Katherine Domb, Aviva Katz).

**Keywords:** DNA methylation, CG methylation, non-CG methylation, CHG methylation, CHH methylation, transposon silencing, gene expression, deamination

## Abstract

DNA methylation in plants occurs in CG, CHG, and CHH sites. While depletion of CG methylation in transposons is associated with ample transcriptional activation, it was mainly studied in species with limited non-CG methylation that is linked to CG methylation. Here we profiled transcription in the moss plant, *Physcomitrella patens*, that has robust non-CG methylation with similar symmetrical CG and CHG methylation levels. Separated contextual methylation mechanisms in *Physcomitrella patens* enabled generation of numerous context-specific hypomethylated mutants. Our transcriptome data show that specific elimination of CG methylation is fully complemented by non-CG methylation. Conversely, exclusive removal of non-CG methylation massively dysregulated genes and transposons. Moreover, CHG methylation silenced transposons stronger than CG methylation. Lastly, we found non-CG methylation as crucial for silencing CG-depleted transposons. These results demonstrate the potency of non-CG methylation in genome regulation and suggest that it evolved due to moderate silencing and/or rapid mutability of methylated CGs.

## Introduction

Cytosine methylation is a prominent DNA modification in many eukaryotes, including plants, animals, and fungi^1–6^. It is vital for regulating various genome activities, including gene expression and transposon silencing^7–11^. One feature of methylation common amongst eukaryotic species, is its preferential targeting to CG sites^1–3,6^. CG methylation can be carried out by various types of DNA methyltransferases (DNMTs). Two major CG methylases are DNMT1 (named MET1 in plants) and DNMT5, which can be found either separately or together in various eukaryotic lineages^3^. The abundance of CG methylation in eukaryotes and its mediation by conserved DNMTs suggest that it is an ancient methylation context in eukaryotes^3^. Besides CG, methylation can also be targeted to non-CG DNA contexts. In mammals, non-CG methylation is enriched in particular tissues and developmental stages, such as neurons and embryonic stem cells^12,13^. In contrast, non-CG methylation in plants is more common and can be found across all tissues and throughout development^1,2,6,14^.

A dominant feature of CG sites is their symmetry at the level of double stranded DNA. Such symmetry is used for maintaining methylation upon DNA replication. For example, DNMT1 together with its cofactor UHRF1 (VIM in plants) were shown to preferentially methylate hemi-methylated CGs, i.e. CG sites methylated on one strand^9,15^. Accordingly, cytosine methylation at CG sites is usually highly correlated between cytosines on both strands and maintained at high efficiency (>80%)^6^. High fidelity of methylation is crucial to maintain the signal through many cell divisions, which is probably why symmetrical CG methylation in eukaryotes is highly conserved between species and abundant within their genomes.

Non-CG methylation can also be oriented in a symmetrical sequence. For example, in plants non-CG methylation occurs at asymmetric-CHH as well as on symmetric-CHG sequence contexts (H= A, C, or T)^1,2,6^. CHG methylation is preferentially catalyzed by chromomethylases (CMTs), which are specialized plant DNMTs that are close homologs of DNMT1^3,16,17^. CMTs are attached to the chromatin by binding to histone H3 Lysine 9-methylated (H3K9me)^18^. While in some plant species, such as *Arabidopsis thaliana (Arabidopsis)*, CHG methylation is inferior to CG methylation, in others (e.g. maize and *Physcomitrella Patens*) CHG methylation can be as robust as CG methylation^6^, suggesting that epigenetic inheritance of non-CG methylation during cell division could be as efficient as CG methylation.

At a functional level, the role of non-CG methylation may be distinct from that of CG methylation. For example, in seed plants CG methylation is usually targeted to genes and transposons, whereas non-CG methylation is enriched mainly in transposons, thus allowing methylation to function distinctively within active and silenced chromatin states^8,9^. In animals, CGs are usually methylated throughout development, whereas non-CGs can function within particular tissues or developmental stages, as in embryonic stem cells and neurons^8,9,13,19^. Non-CG methylation can also supplement or enhance the function of CG methylation. For example, in the diatom *E. huxleyi*, CHG methylation complements CG methylation at linker DNA, where it presumably assists with the positioning of nucleosomes^3^. In plants, both CG and non-CG methylation participate in the silencing of TEs^8,9^. However, thus far transcriptional control by DNA methylation was studied mainly in organisms with distinct CG and CHG methylation efficiencies and cross-interactions between CG and non-CG methylation. Consequently, the particular transcriptional roles of CG and non-CG methylation remains elusive.

On one hand, the long evolution and robust methylation of CG sites in eukaryotes could lead to the development of essential functional readout mechanisms that are more efficient than those of the less conserved non-CG methylation. This hypothesis is supported by the existence of conserved proteins that specifically bind methylated-CG sites in plants and animals^8,9,20^. Conversely, it is possible that newer non-CG methylation evolved to supplement the inefficient silencing mechanism of CG methylation that could have been reduced or bypassed during the arms race with transposons. Thus, resolving the precise functional roles of CG and non-CG methylation is fundamental for understanding the molecular mechanisms and evolution of (contextual) DNA methylation in eukaryotes. Here we investigated the transcriptional roles of CG and non-CG methylation in the moss plant, *P. patens*, which has similar levels and patterns of CG and CHG methylation, limited crosstalk between CG and non-CG methylation, and tolerates the complete loss of various context-specific methylomes. Using this methylation system we were able to unravel the articular functional potency of CG, CHG, and CHH methylation. In contrast to the predominance of CG methylation in eukaryotes, and to previous findings in plant species biased toward CG methylation, our data show that non-CG methylation is superior to CG methylation in silencing TEs and in gene regulation.

## Results

### TEs are strongly controlled by CG methylation in plants with constrained non-CG methylation that is influenced by CG methylation

CG hypomethylation mutants in different plant species caused transcriptional activation of TEs^21–24^. In *Arabidopsis*, CG hypomethylated mutants lead to a broader and stronger upregulation of TE expression than a null non-CG methylation mutant (Figs. 1a-b,j and Supplementary Fig. 1a). While these results imply that MET1 (the CG methylase) is more crucial to silencing TEs than the non-CG methylases, DRM1, DRM2, CMT2, and CMT3 altogether, they do not necessarily mean that CG methylation is more efficient than non-CG methylation in silencing TEs. First, the level of CG methylation in *Arabidopsis*, as well as in other examined plants, is substantially higher than non-CG methylation (Fig. 1c), including within *met1*-upregulated TEs (Supplementary Fig. 1c). Hence, the relatively greater TE activation in *Arabidopsis met1* plants in comparison to *drm1drm2cmt2cmt3* (*ddcc*) could be related to the robustness of CG methylation versus the limited non-CG methylation signal. Second, in all examined species, CG hypomethylation in *met1*-upregulated TEs is associated with considerable reduction in non-CG methylation (Figs. 1d-f), which in *Arabidopsis* are localized preferentially to TE edges (Figs. 1g-h) or spread out throughout the entire element (Figs. 1i-j). Thus, the activation of TEs in these plants cannot be associated only to CG hypomethylation.

**Fig. 1.**
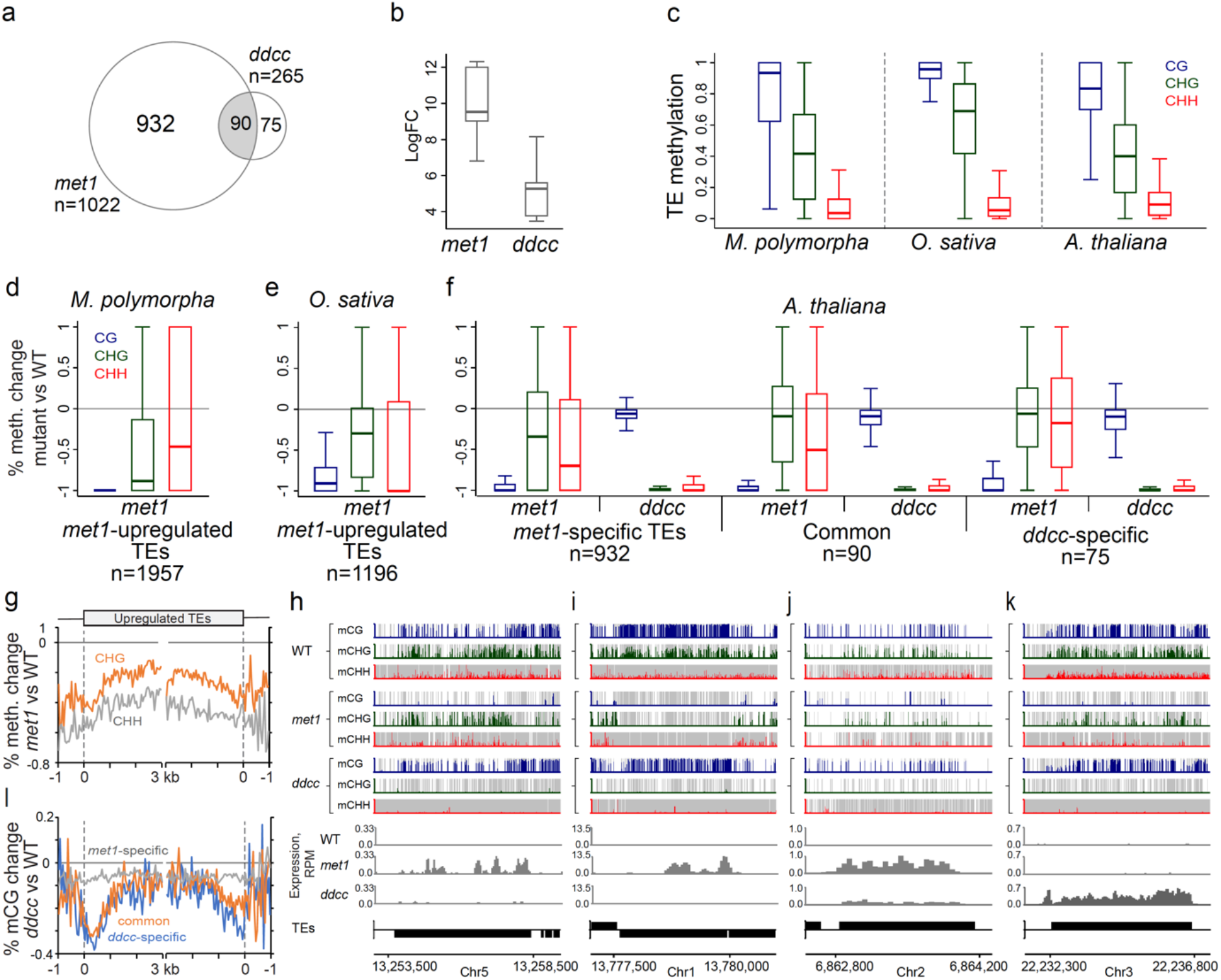
TEs are strongly controlled by CG methylation in plants with constrained non-CG methylation that is influential by CG methylation. (a) Venn diagram of upregulated TEs in *met1*^32^ and *ddcc*^51^ mutants. (b) Box plots showing RNA log fold change (logFC) of TEs upregulated in both *met1* and *ddcc* mutants. (c) Box plots of WT DNA methylation in methylated TEs separated to CG, CHG, and CHH contexts in *M. polymorpha*^22^, *O. sativa*^23^, and *A. thaliana*^32^ methylated TEs. Methylated TEs have at least 5% averaged total DNA methylation. Raw BS-seq reads of published and current datasets were aligned, quality controlled, and quantified for methylation using the same computational pipeline as described in the methods section. (d-e) Box plots of percent-methylation-change between averaged methylation (CG, CHG, or CHH) in WT versus mutant calculated for 50 bp windows within upregulated TEs in *M. polymorpha* (d) and rice (d). (f) Box plots of percent-methylation-change calculated as in (d-e) within TEs commonly and exclusively upregulated in *A. thaliana met1* and *ddcc* mutants. (g) Patterns of percent-methylation-change in TEs upregulated in the *A. thaliana met1* mutant. *met1*-upregulated TEs were aligned at the 5′ or 3’ ends, and CHG or CHH methylation-change between *met1* and WT were averaged within 50 bp intervals. The dashed lines represent points of alignment. (l) Patterns of percent-methylation-change along TEs commonly or exclusively upregulated in *A. thaliana met1* and *ddcc* mutants. *met1*- and/or *ddcc*-upregulated TEs were aligned as in (g) and CHG or CHH methylation-change between *ddcc* and WT were averaged within 50 bp intervals. (h-k) Genome browser snapshots of upregulated-TEs in *A. thaliana met1* and *ddcc* mutants. *met1*-specific upregulated TEs (h-i), commonly upregulated-TEs in *met1* and *ddcc* (j), *ddcc*-specific upregulated TEs (k). CG, CHG and CHH methylation levels in 10-bp windows resolution are shown in blue, green and red, respectively. Grey bars represent the presence of covered sites in each methylation context in the given 10-bp window. Expression is shown as normalized reads coverage (RPM) in 50 bp windows. TEs oriented 5′ to 3′ and 3′ to 5′ are shown above and below the line, respectively.

The presence of *ddcc*-specific activated TEs could have suggested that at some TEs, non-CG methylation and CG methylation are crucial and trivial for silencing, respectively (Fig. 1k). However, analysis of CG methylation in *ddcc* across the sequence of different activated TE subgroups, revealed it to be particularly depleted at the edges (primarily 5-prime ones) of only *ddcc*-activated TEs and not in *met1*-specifically activated ones (Figs. 1k-l). Thus, we cannot exclude the possible contribution of CG hypomethylation to the activation of TEs in the null non-CG methylation mutant, *ddcc*.

To summarize, while *Arabidopsis* provides us viable mutants with near-complete hypomethylation in either CG and non-CG contexts, the difference in CG and non-CG methylation efficiencies and the mixed-context hypomethylation effect in each of the mutants challenge our ability to differentiate between the precise transcriptional roles of CG and non-CG methylation.

### Total and specific elimination of CG methylation in *P. patens* does not activate TE expression

Here we investigated the functional roles of CG and non-CG methylation in *P. patens*, which has a robust CHG methylation level that is similar to that of CG (~80%) [^1,17,25^; Fig. 2a]. Additionally, *P. patens* has three DNMTs, MET, CMT, and DNMT3b, which when compromised generate viable CG, CHG, and CHH null methylation mutants, respectively [^17,26,27^; Fig. 2a]. Finally, a small subset (120) of transcribed hypomethylated TEs in wild type (Supplementary Figs. 2a-c), suggests that *P. patens* TE expression is controlled by methylation.

**Fig 2.**
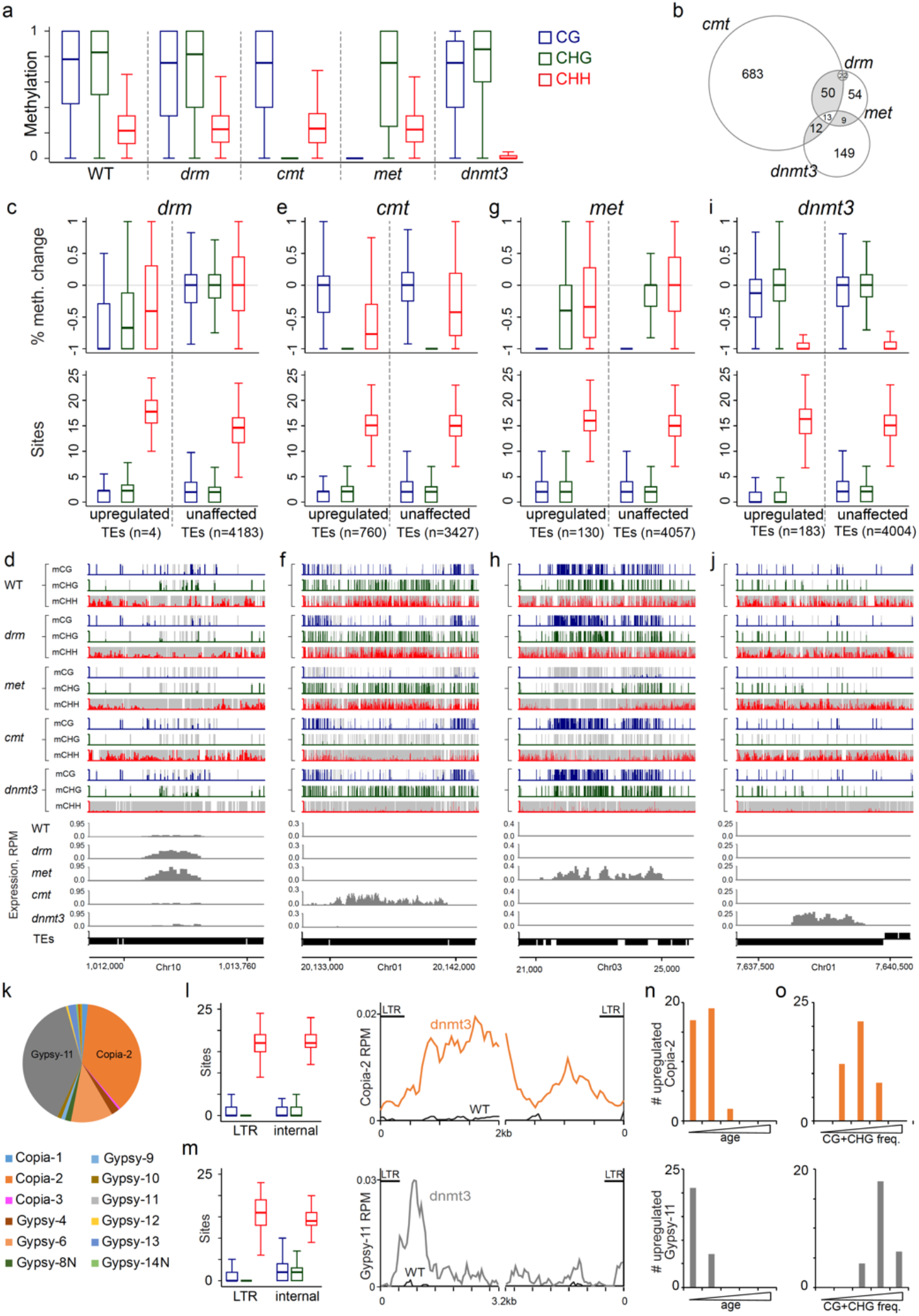
CG methylation is complemented by non-CG methylation and CHH methylation silences CG and CHG depleted TEs. (a) Box plots of CG, CHG and CHH methylation in methylated TEs quantified in 50-bp windows in *P. patens* WT and indicated mutants. *drm* is mutated in DRM1 and DRM2. *dnmt3* is mutated in DNMT3a and DNMT3b. (b) Venn diagram of upregulated TEs in *met*, *cmt*, *dnmt3* and *drm* mutants. (c,e,g,i) – Box plots of percent-methylation-change (top panel) and CG, CHG and CHH sites frequency (bottom panel) in 50-bp windows within upregulated or unaffected TEs in each mutant. Unaffected TEs considered as ones that were expressed in all mutants in this study (single and doubles) except for the indicated mutant. (c) *drm*. (e). *cmt*; note the CHH loss and lower total CG sites frequency in upregulated TEs. (g) *met*; note the non-CG methylation loss in upregulated TEs; (i). *dnmt3*; note the low CG and CHG sites density in upregulated TEs. (d,f,h,j) Genome browser snapshots of TEs upregulated in *drm* (d), *cmt* (f), *met* (h) and *dnmt3* (j) mutants. (k) Pie chart of TE families upregulated in *dnmt3* mutant. (l) Box plots of CG, CHG and CHH sites frequency in 50 bp windows within LTRs and internal sequences of *dnmt3*-upregulated intact Copia-2 (left), and mRNA-seq reads distribution along *dnmt3* upregulated Copia-2. (m) Same as in L just for upregulated intact Gypsy-11 in *dnmt3*. (n) Bar graph of the number of *dnmt3-*upregulated intact Copia-2 (top panel) and Gypsy-11 (bottom panel) over five age quantiles. (o) Bar graph of the number of *dnmt3-*upregulated intact Copia-2 (top panel) and Gypsy-11 (bottom panel) over five quantiles of symmetric (CG and CHG) sites frequency.

To investigate the effect of DNA methylation on TE expression in *P. patens*, we profiled transcriptomes of the following methylation mutants, *met*, *cmt*, *dnmt3b* (termed *dnmt3*), and the double mutant *drm1drm2* (termed *drm*). Consistent with their substantial hypomethylation phenotypes (^17^; Fig. 2a), *cmt*, *dnmt3*, and *met* mutants caused a considerable activation of TEs (130-760 TEs), whereas *drm* in correlation to its minute hypomethylation effect activated only four TEs (Figs. 2a-b). Upregulated TEs in *cmt*, *dnmt3*, and *met*, were only partially overlapping, where *met* had the lowest activation effect (130 TEs), *dnmt3* showed a slightly higher effect (183 TEs), and *cmt* had the greatest TE activation response (760 TEs) (Fig. 2b). Overall, these results suggest that DNA methylation restricts TE activity in *P. patens*, where each DNMT controls a distinct subset of TEs, and that CMT has the strongest effect among all PpDNMTs.

To better understand the role of CG and non-CG methylation on TE regulation, we next analyzed the CG, CHG and CHH frequency and methylation level within upregulated TEs in each of the mutants. Firstly, we found that *drm*-upregulated TEs were already partially methylated and expressed in wild type (Supplementary Fig. 2d). Hypomethylation and expression were enhanced in *drm* (Figs. 2c-d). This finding is consistent with PpDRMs preference to methylate actively transcribed TEs^17^. Secondly, we found that in *cmt* a total CHG hypomethylation (98.4%) is associated with substantial CHH hypomethylation (57%) within *cmt*-upregulated TEs (Figs. 2e-f), suggesting that TE activation in *cmt* cannot be attributed solely to the loss in CHG methylation (Fig. 2f). Thirdly, we discovered that in comparison to the general CG-specific hypomethylation effect observed in *met* (Fig. 2a), within its 130 upregulated TEs, methylation is reduced also at non-CG sites by about 40% (Figs. 2g-h). These results suggest that except for a small number of TEs that were hypomethylated in all methylation contexts, a total and specific elimination of CG methylation in *met* did not affect TE expression, suggesting for functional complementation by non-CG methylation.

### CHH methylation is crucial for silencing CG and CHG depleted TEs

In contrast to *cmt* and *met* mutants that lost substantial amount of methylation in multiple methylation contexts at upregulated TEs, in the *dnmt3* mutant, methylation in upregulated TEs was lost at CHH sites but remained mostly intact in CG or CHG sites (Figs. 2i-j). Interestingly, we discovered that *dnmt3*-upregulated TEs are substantially depleted of CG and CHG sites (Figs. 2i-j). Upregulated TEs in *dnmt3* were composed of mainly two types of long terminal repeat (LTR) elements, Gypsy-11 and Copia-2 (Fig. 2k). In *dnmt3*-upregulated Copia-2, CG and CHG sites were depleted across the entire elements (Fig. 2l). In comparison, CG and CHG sites in *dnmt3*-upregulated Gypsy-11 were particularly depleted within LTR regions (Fig. 2m). In concordance with their CG/CHG site depletion profiles, Copia-2s were transcribed in *dnmt3* throughout their entire sequences whereas Gypsy-11s were preferentially transcribed in proximity to the LTR sequences (Figs. 2l-m). Additionally, our data show that the TE-upregulation in *dnmt3* is enriched among young and moderately CG/CHG-depleted TEs (Figs. 2n-o). Cytosine methylation is known to be mutagenized by deamination^28^. By quantifying the frequency of symmetric and asymmetric sites over TE age, we found CG and CHG sites to have a wider range than CHH sites, i.e. ~60% and ~8% change in frequency for symmetric and asymmetric sites, respectively (Supplementary Figs. 3a-b), suggesting a faster mutability rate for symmetric versus asymmetric sites. Together, these results suggest that CHH methylation is crucial for the silencing of a transient phase in the life of TEs while being depleted of symmetrically (methylated) cytosine sites.

**Fig. 3.**
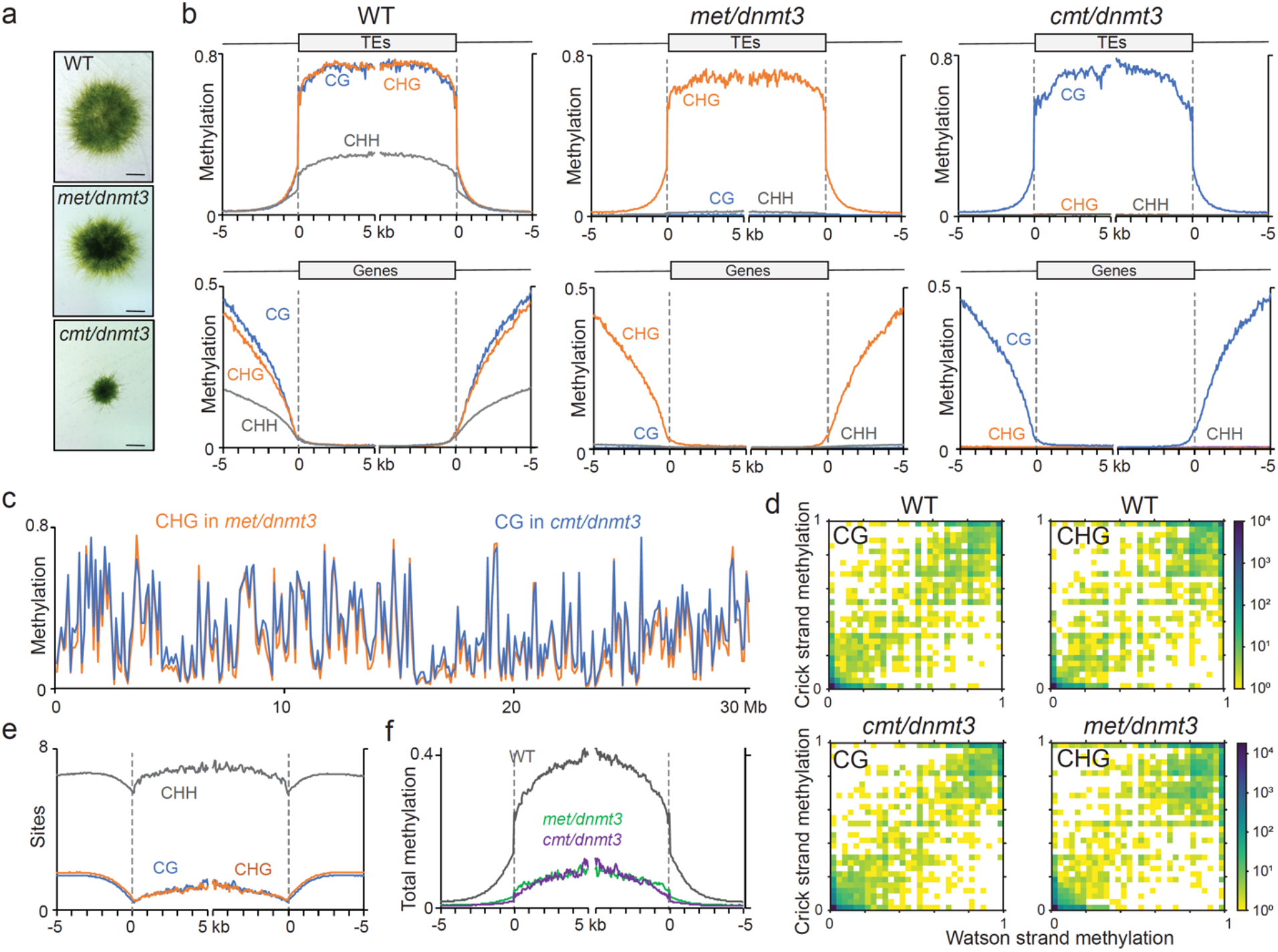
Generating comparable CG- and CHG-specifically methylated *P. patens* epigenomes. (a) Pictures of representative WT and mutant plants at the gametophytic stage. Statistical analysis of the phenotype can be found in Supplementary Fig. 4a. (b) Patterns of CG, CHG and CHH methylation in WT and *met/dnmt3* and *cmt/dnmt3* mutants, along TEs (top) and genes (bottom). (c) Average CG methylation in *cmt/dnmt3* and CHG methylation in *met dnmt3* along *P. patens* chromosome 1 averaged in 100-kb sliding window. (d) Density scatter plot correlating CG and CHG methylation within same sites on each of the strands. (e) Patterns of CG, CHG and CHH sites frequency in P. patens TEs. (f) Patterns of total DNA methylation in TEs in WT and in *met/dnmt3* and *cmt/dnmt3* mutants.

### Generating comparable CG- and CHG-specifically methylated epigenomes

Upregulated TEs in wild type or the single mutants, *met*, *cmt*, and *dnmt3* were typically hypomethylated in multiple contexts, i.e. CG and/or CHG and CHH, or hypomethylated in CHH and depleted of CG and CHG sites (Figs. 2c-i and Supplementary Figs. 2b-c). These results suggest a functional complementation between symmetrical and asymmetrical methylation in *P. patens*. To test this hypothesis and to explore the greater potential of DNA methylation in TE regulation in *P. patens*, we next aimed at generating mutants depleted of methylation in multiple contexts. We succeeded at generating viable *cmt*/*dnmt3* and *met*/*dnmt3* double mutants (Fig. 3a and Supplementary Fig. 4). However, following numerous attempts we failed to generate a *met*/*cmt* double mutant. Neither directed mutagenesis of knocking out MET on *cmt* background nor mutating CMT on *met* background, yielded a *met*/*cmt* double mutant, suggesting that a complete loss of CG and CHG methylation is lethal. Methylome analysis of *cmt*/*dnmt3* found it to be entirely depleted of CHG and CHH methylation (Fig. 3b). In comparison, *met*/*dnmt3* methylome was entirely depleted from CG and CHH methylation (Fig. 3b). Consequently, *met*/*dnmt3* and *cmt*/*dnmt3* were CHG- and CG-specifically methylated, respectively. Notably, CG and CHG methylation profiles in genes, TEs, and along chromosomes, in *cmt*/*dnmt3* and *met*/*dnmt3*, respectively, were similar and close to their normal levels in wild type (Figs. 3b-c). Furthermore, both CG and CHG methylation remained symmetrical, i.e. similarly methylated on both strands (Fig. 3d). Finally, as CG and CHG frequencies are generally similar in *P. patens* TEs (Fig. 3e), total methylation in both mutants is comparable (Fig. 3f).

**Fig. 4.**
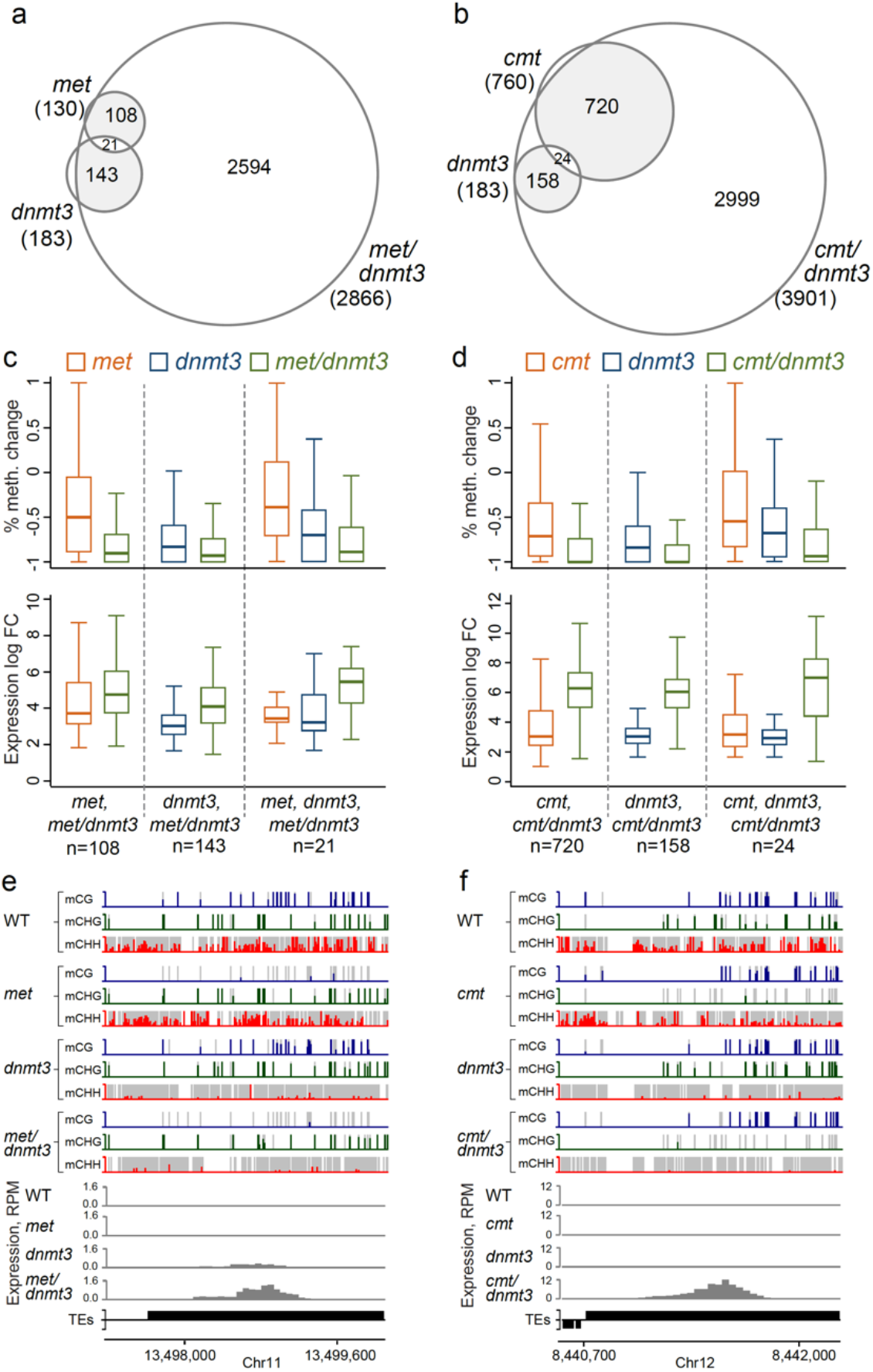
Synergistic transposon silencing by symmetric and asymmetric methylation. (a-b) Venn diagrams of upregulated TEs in single *(dnmt3* and *met* or *cmt*) and double (*met/dnmt3* or *cmt/dnmt3*) mutants. (c-d) Box plots of percent of total methylation change between WT and indicated mutants (top) and RNA log fold change (bottom) of TEs commonly upregulated in single and double mutants. (e-f). Genome browser snapshots illustrating single examples of synergistically upregulated TEs in the double mutants.

To summarize, the close-to-normal and specific CG methylation in *cmt*/*dnmt3* (Figs. 3a-b) suggests a weak feedback of non-CG (CHG and CHH) methylation on CG methylation. Similarly, close-to-normal and specific CHG methylation in *met*/*dnmt3* (Figs. 3a-b) suggests a weak influence of CHH and CG methylation on CHG methylation. Finally, and most importantly, we succeeded in generating two distinct epigenomes consisted with either CG-specific or CHG-specific methylation at comparable levels, frequencies, and chromosomal distributions (Fig. 3). These unique epigenomes, can be utilized for studying the interplay between symmetrical (CG/CHG) and asymmetrical (CHH) methylation as well as between the two symmetrical contexts, CG and CHG.

### Functional complementation between symmetric (CG/CHG) and asymmetric (CHH) methylation in silencing TEs

To test for functional complementation between symmetric and asymmetric methylation we next profiled transcriptomes and analyzed TE regulation in the double mutants, *met*/*dnmt3* and *cmt*/*dnmt3* (Fig. 4). In correspondence with their strong hypomethylation profiles (~80% reduction in total methylation), both double mutants had a substantially stronger activation of TEs in comparison to their single mutants (Figs. 4a-b). In *met*/*dnmt3* mutant, 16 and 22 times more TEs were activated in comparison to single *dnmt3* and *met* mutants, respectively (Fig. 4a). In *cmt*/*dnmt3* mutant, 21 and 5 times more TEs were activated in comparison to *dnmt3* and *cmt* single mutants, respectively (Fig. 4b). Additionally, most upregulated TEs in the single mutants (>90%) were found to be upregulated in the relevant double mutants (Figs. 4a-b). Lastly, all groups of activated TEs in the single mutants, i.e. which were activated in either or both single mutants, showed enhanced activation in the relevant double mutant, which was associated with greater hypomethylation (Figs. 4c-f). Overall, these synergistic silencing effects indicates a strong functional-complementation between symmetrical CG and CHG methylation to that of asymmetric CHH methylation in controlling TE expression.

### CHG methylation is superior to CG methylation in silencing TEs

The presence of thousands of upregulated TEs in *cmt*/*dnmt3* and *met*/*dnmt3* suggests that each of the symmetrical methylated contexts, CG and CHG, by themselves are not sufficient to silence a substantial number of TEs in *P. patens* genome. At the same time, these double mutants provide us the unique opportunity to analyze the transcriptional alterations in comparable methylomes that are either totally depleted of CG and CHH methylation or from CHG and CHH methylation, thus allowing us to investigate the strength of the remaining comparable CG and CHG methylation on genome regulation.

Our differential expression analysis found more upregulated TEs in *cmt*/*dnmt3* (3901) than in *met*/*dnmt3* (2866) (Fig. 5a). Additionally, among commonly upregulated TEs, i.e. activated in both mutants, expression in *cmt*/*dnmt3* was substantially higher (~5x) than in *met*/*dnmt3* (Fig. 5b). To verify the role of methylation in TE activation, we profiled methylation level and frequency in various upregulated subgroups (Figs. 5c-h). Among *met*/*dnmt3*-specifically activated TEs (n=286), frequency of CG sites was higher than that of CHG (Fig. 5c). Additionally, *met*/*dnmt3*-specifically activated TEs were substantially hypomethylated at CHG sites in *met*/*dnmt3*. Together, these results suggest that reduction in CHG methylation and its frequency resulted in the upregulation of *met*/*dnmt3*-specific TEs. Similarly, in *cmt*/*dnmt3*-specifically upregulated TEs (n=1321) the CG content was exceptionally low, causing a near-complete hypomethylation in all methylation contexts in *cmt*/*dnmt3* (Figs. 5d,g). Consequently, one reason for the greater role of CHG methylation over that of CG methylation in TE regulation is the depletion of CG sites among a substantial number of TEs. Within commonly-activated TEs (n=2580), CG and CHG frequencies and methylation levels are similar, with a slightly higher level of CG methylation in *cmt*/*dnmt3* than CHG methylation in *met*/*dnmt3* (Figs. 5e,h). Thus, the stronger expression of commonly-activated TEs in *cmt*/*dnmt3* versus *met*/*dnmt3* (Figs. 5b,e,h), implies that CG methylation is less efficient than CHG methylation in silencing TEs. We next checked for localized methylation and expression patterns across intact TEs in the four main TE family types that were commonly activated in the double mutants, i.e. Gypsy-11, Copia-2, Gypsy-4, and Gypsy-6 (Supplementary Fig. 5a). In Gypsy-4, expression is limited to the 5’ LTR sequence and is substantially stronger in *cmt*/*dnmt3* than in *met*/*dnmt3* (Supplementary Fig. 5b). Gypsy-4 LTR expression in *cmt* and *cmt*/*dnmt3* mutants is associated with a strong localized CG-hypomethylation (Supplementary Fig. 5b). These results imply that CMT can regulate CG methylation at particular TE sequences, and that reduction of methylation in all three sequence contexts caused the stronger transcription of Gypsy-4 LTRs in *cmt*/*dnmt3* over *met/dnmt3*. In Gypsy-11, which composed the majority of commonly-upregulated TEs (Supplementary Fig. 5a), transcription in *cmt/dnmt3* is over 5-fold stronger than in *met/dnmt3* and initiates within the 5’-LTR and decreases at about 1kb downstream from it (Fig. 5i). CG and CHG methylation levels are similar around and downstream to Gypsy-11 transcription start site (TSS) (Figs. 5h,i). In comparison, CG and CHG frequencies are increased and decreased, respectively, upstream to Gypsy-11 TSS (Fig. 5i). Similarly to Gypsy-11, CG and CHG sites are enriched and depleted, respectively, in the LTR regions of Copia-2 elements, while CG and CHG methylation remained normal and similar in *cmt*/*dnmt3* and *met*/*dnmt3*, respectively (Supplementary Fig. 5c). CHH methylation in Gypsy-11 and Copia-2 is similarly depleted in both double mutants (Fig. 5i and Supplementary Fig. 5c). Hypomethylation of CG islands around TSS are commonly associated with active transcription^29,30^. Therefore, the stronger Gypsy-11 and Copia-2 activation in *cmt*/*dnmt3* over *met*/*dnmt3*, despite the remains of CG methylation around their TSS, corroborate the trivial role of CG methylation and the strength of CHG methylation in the transcriptional regulation of such TEs. Gypsy-6 is another group of TEs that were activated in both double mutants (Supplementary Fig. 5a). Gypsy-6 are hypomethylated and expressed at their LTR regions already in wild type (Fig. 5j). While CG and CHG frequency and methylation are comparable across Gypsy-6 in *cmt*/*dnmt3* and *met*/*dnmt3*, respectively, in *met*/*dnmt3* expression is focused to the LTR regions, whereas in *cmt*/*dnmt3* expression is increased across the entire transposon (Fig. 5j). Altogether, Gypsy-11, Copia-2, and Gypsy-6 expression and methylation patterns in the double mutants suggest that CHG methylation is superior to CG methylation in silencing transposons.

**Fig. 5.**
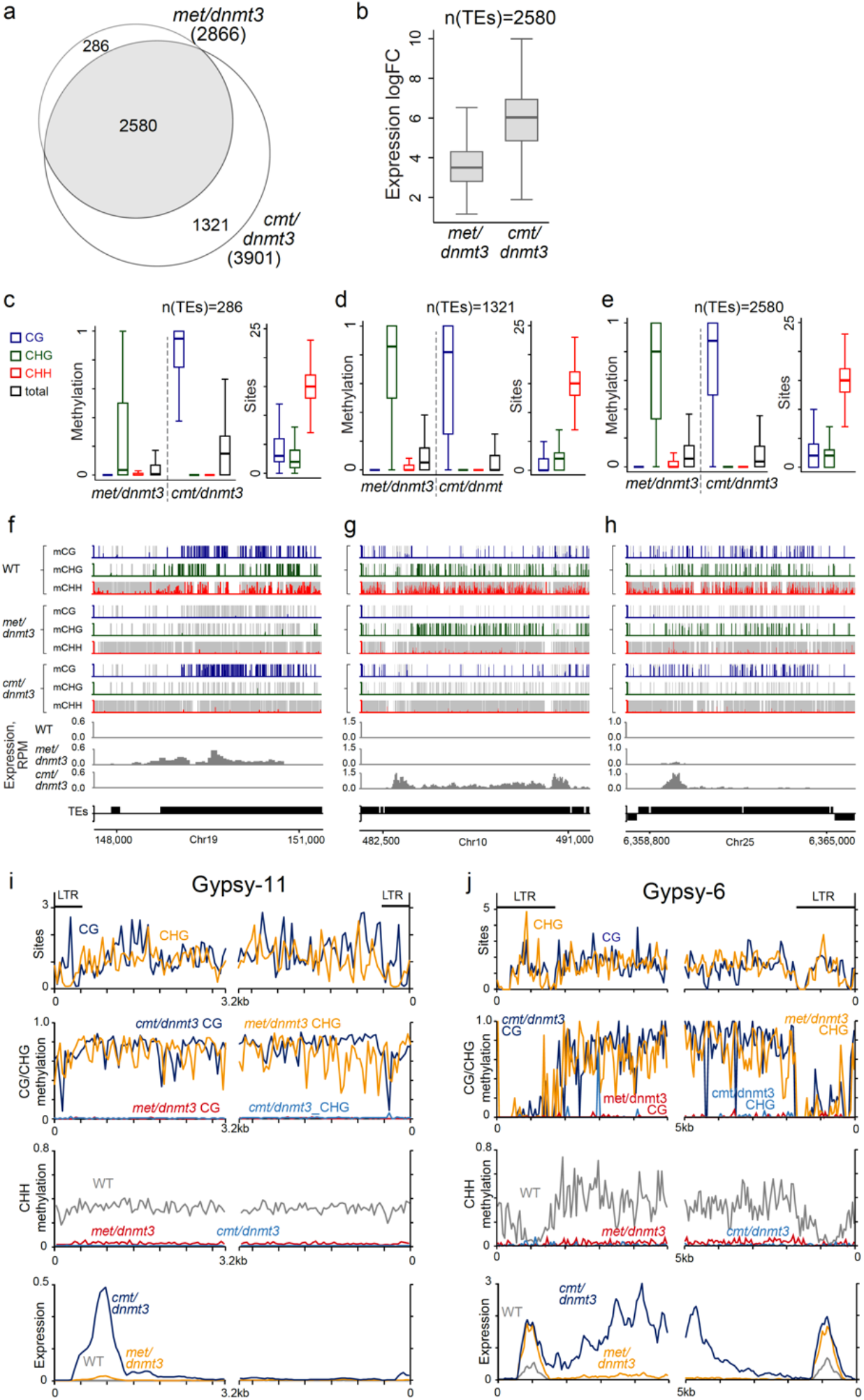
CHG methylation is superior to CG methylation in silencing TEs. (a) Venn diagram of upregulated TEs in *met/dnmt3* and *cmt/dnmt3* mutants. (b) Box plots of expression of TEs upregulated in both *met dnmt3* and *cmt dnmt3* mutants. (c-e) Box plots of CG, CHG, and CHH methylation level and site frequency in 50 bp windows within TEs upregulated exclusively in either *met/dnmt3* mutant (c), *cmt/dnmt3* mutant (d), or in both double mutants (e). (f-h) Genome browser snapshots of TEs upregulated exclusively in *met dnmt3* mutant (f), *cmt dnmt3* mutant (g), and upregulated in both double mutants (h). CG, CHG, and CHH methylation are in blue, green, and red, respectively. Normalized RNA reads level are in gray. (i, j) Patterns of CG and CHG sites distribution (first panel), CG or CHG methylation in *met/dnmt3* and *cmt/dnmt3* mutants (second panel), CHH methylation in WT and double mutants (third panel), and expression (RPM) (forth panel) averaged in 50 bp windows, along intact upregulated Gypsy-11 (i) and Gypsy-6 (j) TEs in the double mutants (n=2580). Intact TEs were aligned at the 5’ or the 3’ ends (zero at the x-axis). LTR sequences are marked in the top panel.

### Complementation and superiority of non-CG methylation over CG methylation in gene regulation

While DNA methylation in *P. patens* is mostly targeted to TEs, transposon methylation could influence the expression of nearby genes^31,32^. In comparison to TEs, where internal methylation auto-silencing the elements, in genes methylation commonly regulates expression (activation and suppression) by targeting external-genic elements^33–38^. Additionally, genetic interaction networks are more robust in genes than in transposons. Accordingly, it is expected that mutants defected in methylation will have complex differential gene expression profiles composed of up and down regulations which are directly and indirectly influenced by DNA hypomethylation.

Our data show that the number of differentially expressed genes are correlated to the number of activated transposons in the various methylation mutants (Fig. 6a). In correlation to its trivial methylation and TE activation effects, no genes were differentially expressed in the *drm* mutant (Fig. 6b). In all other than *drm* mutants, we detected both activated as well as suppressed genes (Fig. 6b). A significantly higher linear correlation was found between the number of activated TEs and the number of activated genes (r=0.94, p<0.006) versus the number of suppressed genes (r=0.79, p<0.06) (Fig. 6c). Most upregulated genes were not methylated internally (Fig. 6d). In comparison, we found adjacent intergenic regions, especially promoters, of upregulated genes to be more methylated in wild type than intergenic regions of unaffected genes (Fig. 6e) and to be enriched with DMRs in the mutants (Fig. 6f). Conversely, promoter regions of suppressed genes had mostly lower methylation in wild type than the background of unaffected genes (Fig. 6e). Interestingly, we found higher frequency of activated genes located in close proximity (within 10 kb) of activated TEs (Fig. 6g). Despite the elevated proximity to expressed TEs, most upregulated genes were not the result of continuous expression from adjacent TEs (Fig. 6d). Overall, these results suggest that TE methylation in *P. patens* controls gene expression.

**Fig. 6.**
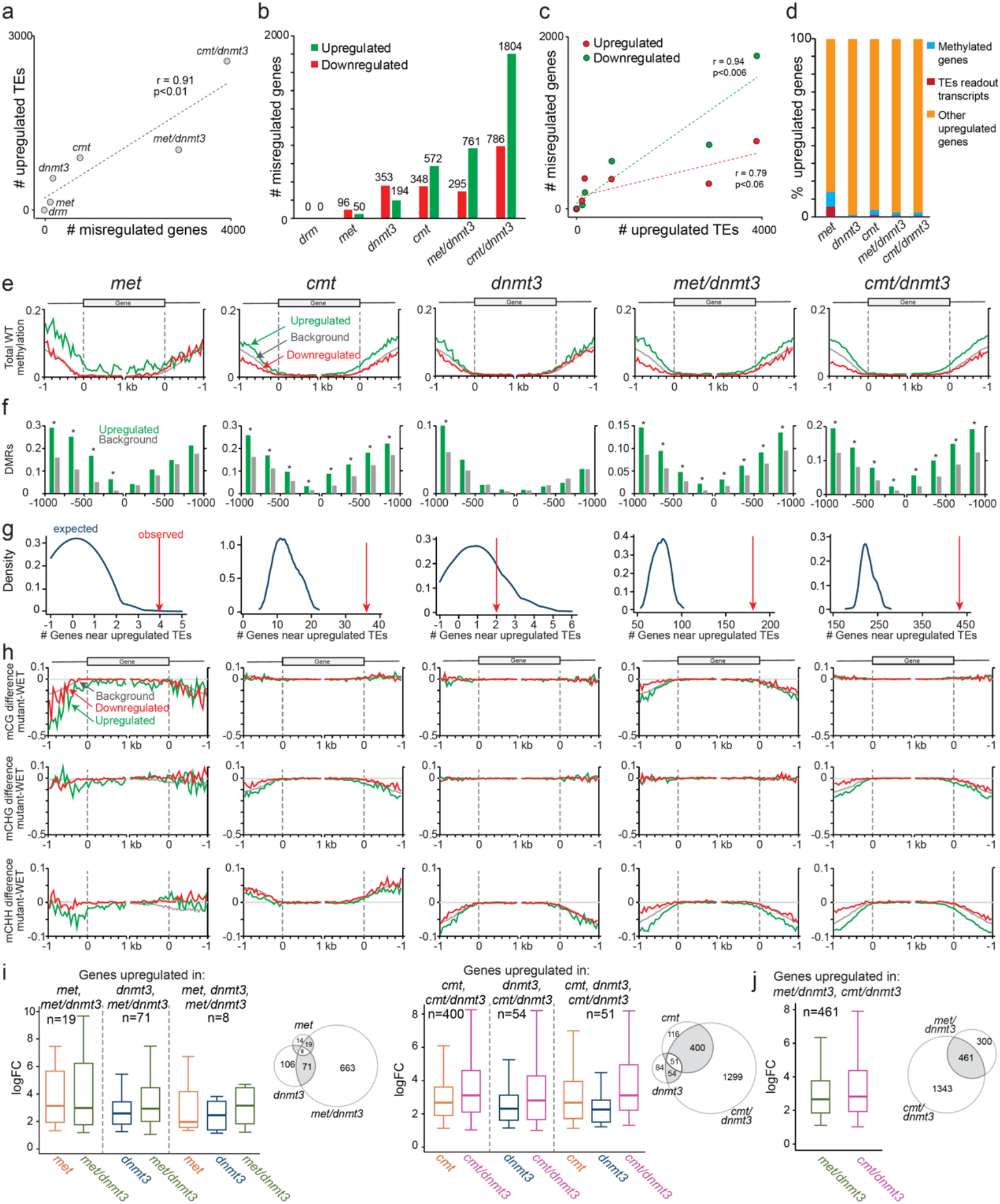
Complementation and superiority of non-CG methylation over CG methylation in gene regulation. (a) Scatter plot between upregulated TEs and differentially expressed genes in indicated mutants. (b) Number of significantly upregulated and downregulated genes in the indicated mutants. (c) Scatter plot between upregulated TEs and upregulated (green) or downregulated (red) genes in single and double mutants. (d) Percentage of methylated genes (>5% methylation in non-CG) or ones that are continuously expressed from an adjacent TE, out of the total upregulated genes. (e) Pattern of total DNA methylation in WT of upregulated, downregulated or unaffected (background) genes in indicated mutants. Genes were aligned at the TSS and TTS (dashed lines) and methylation averaged in 50 bp windows. (f) Frequency of upregulated and unaffected genes with a hypo-DMR in the indicated mutants (as in e) within 1 kb away from TSS or TTS. (g) Density plot of randomly expected upregulated genes within 10 kb of upregulated TE in the indicated mutants (as in e). Red arrows represent the observed overlap between upregulated genes and TEs. Random distribution plots include 100 runs on an equal number of genes retrieved randomly from expressed genes. (h) Pattern CG, CHG and CHH methylation difference between indicated mutants (as marked in e) and WT. Genes were separated to upregulated, downregulated or unaffected (background) genes in the indicated mutants. Arrows mark non-CG hypomethylation upstream to TSS of met-upregulated genes. (i-j) Box plots of expression (LogFC) of genes commonly-upregulated in the indicated mutants as illustrated in the Venn diagrams.

We detected only 50 upregulated genes in the *met* mutant (Fig. 6b). Similarly to the effect in TEs, *met*-upregulated genes have been associated with a loss in non-CG methylation. Promoter regions, the first 500 bp upstream to the TSS, of *met*-upregulated genes were particularly hypomethylated in CHG and CHH methylation (Fig. 6h). Thus, total and specific elimination of CG methylation hardly activated genes in *P. patens*. In comparison to the *met* mutant, *dnmt3*, *cmt*, and *cmt*/*dnmt3* mutants, were depleted mainly in non-CG methylation and hardly in CG methylation (Fig. 6h) but caused to a greater gene activation (194-1804 genes; Fig. 6b), suggesting that non-CG methylation is more crucial than CG methylation to gene regulation.

Double mutants showed a higher number of activated genes than the sum of upregulated genes in single mutants (Fig. 6b). Additionally, differential expression level of genes that were activated in the double and single mutants, was higher in the double mutants (Fig. 6i). While, *cmt*/*dnmt3* and *met*/*dnmt3* show specific and comparable CG and CHG methylation levels respectively, at adjacent intergenic regions (Fig. 3b), *cmt*/*dnmt3* had 2.4 times more upregulated genes than *met*/*dnmt3* (Fig. 6b). Furthermore, among commonly-upregulated genes in the double mutants, in *cmt*/*dnmt3* the level of upregulation was stronger than in *met*/*dnmt3* (Fig. 6j). The prominent differential gene expression in *cmt*/*dnmt3* versus *met*/*dnmt3* is correlated to its altered morphological phenotype (Fig. 3a and Supplementary Fig. 5a).

Overall, these results suggest that similar to TEs, a total and specific elimination of CG methylation hardly affects gene expression and is complemented by non-CG methylation. Our data also show that in *P. patens* symmetric (CG and CHG) and asymmetric (CHH) methylation function synergistically in regulating gene expression, and that CHG methylation is superior to CG methylation in gene regulation.

## Discussion

Previous genetic studies on plant DNA-hypomethylation mutants have shown that CG methylation has a more pronounced role in TE silencing than non-CG methylation^21–24^. These results potentially support the importance of CG methylation and its relative strength over non-CG methylation. However, these studies were mainly conducted in species with more robust CG methylation than non-CG methylation, and with a strong influence of CG methylation on the non-CG methylation (Figs. 1 and 7a), which impede concluding the relative role of each of the contexts per methylated site. Here, we chose to unravel the particular roles of CG and non-CG methylation on gene and TE expression, by investigating it in the moss plant, *P. patens*, which has a robust CHH methylation, a comparable level and distribution of symmetric CHG and CG methylation, as well as a minimal influence of CG methylation on non-CG methylation and vice versa (Figs. 2-3). Furthermore, *P. patens* mutants tolerates complete loss of particular or multiple methylation contexts, including ones eliminated of CHH methylation combined with a complete removal of either CG or CHG methylation. In this system, we found that a specific elimination of CG methylation is functionally complemented by non-CG methylation (Figs. 4e and 7b). In contrast, elimination of non-CG methylation was not complemented by CG methylation (Figs. 4f and 7b). Furthermore, we show that between the two symmetrical methylation contexts, CHG is superior to CG in controlling the expression of TEs and genes (Figs. 5,6 and 7b). Lastly, we found non-CG methylation to be crucial for silencing TEs that are depleted of CG sites (Figs. 2j,5g, and 7c). These findings elucidate the strength and significance of non-CG methylation on genome regulation.

**Fig. 7.**
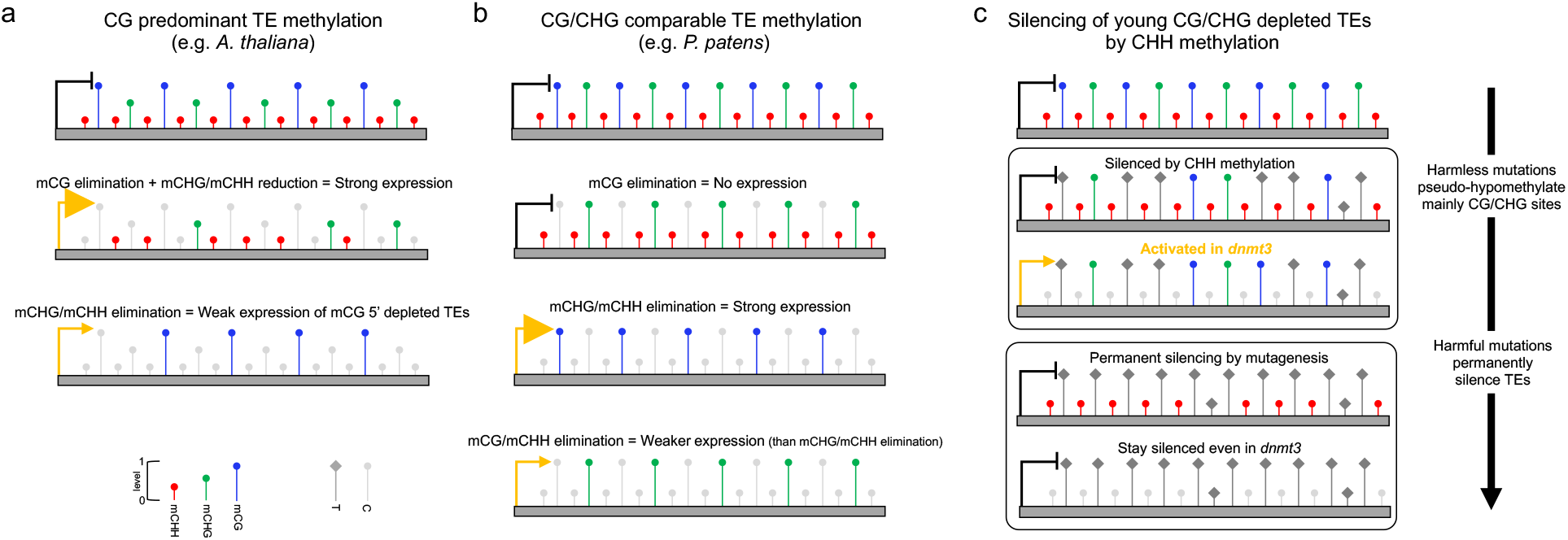
A model for TE transcriptional control by CG, CHG, and CHH methylation in *plants*. (a) Transposon transcriptional control in genome with mCG predominant genome (high mCG and low CHG methylation) with influence on non-CG methylation. Removal of CG methylation and reduction in non-CG methylation cause to a strong transcriptional activation. In contrast, removal of non-CG methylation activate a smaller subset of TEs that are depleted of CG methylation at their 5’. (b) Transposon transcriptional control in genome with comparable levels of mCG and mCHG and with limited crosstalk between CG and non-CG methylation. Total elimination of only CG methylation does not activate TE expression. In contrast, total elimination of non-CG methylation causes to a robust TE expression, suggesting a null silencing effect of mCG. Total elimination of CG and CHH methylation activated TE expression but to a weaker level than total elimination of CHG and CHH, suggesting for functional complementation between CG and CHH methylation and for stronger silencing effect by CHG methylation than by CG methylation. (c) A model for pseudo-hypomethylation by mutagenesis and the role of CHH hypomethylation in protecting pseudo-hypomethylated elements from being activated. Methylated cytosines are converted to thymines via deamination. Highly methylated cytosines, like CG and CHG, are more rapidly deaminated. Harmful conversions of methylated cytosines to thymines eventually lead to permanent silencing of TEs. However, harmless mutations in symmetric sites of relatively young TEs could lead to transcriptional activation upon removal of CHH methylation (e.g. in *dnmt3*).

The conservation and persistence of CG methylation along eukaryotic evolution driven TEs to adapt counteracting mechanisms, such as VANC proteins that induce hypomethylation^36^. Our finding that CHH methylation silences young TEs depleted of symmetric CG and CHG sites (Fig. 2) suggests an alternative mechanism for TEs to avoid methylation by the elimination of potentially methylated sites. Elimination of methylated cytosines happen by deamination leading to the accumulation of C:G to A:T transition mutations^28^. Sufficient harmful mutations would lead to permanent silencing of the element (Fig. 7c). However, insufficient harmless mutations of methylated cytosines could place the host at risk due to pseudo-hypomethylation of the elements (Fig. 7c). Moreover, while increased mutagenesis might eliminate the ability of intact elements to transpose, non-mobile TE fragments and solo-LTRs can still pose a risk to the host by acting as cryptic promoters. Under such circumstances, one could see how more frequent and less efficient methylation mechanism, such as that of CHH, would evolve to protect the host genome from malicious elements during their transient methylation-driven incompetent mutagenesis phase (Fig. 7c). In *P. patens*, both CG and CHG sites are robustly methylated (~80% in TEs). High methylation level is efficient for silencing, but at the same time could promote a more rapid mutagenesis rate than the less efficient CHH methylation signal. Correspondingly, we found symmetric CG and CHG sites in *P. patens* TEs to have a broader frequency range across TE age than the one we found for CHH sites (Supplementary Figs. 3a-b), concluding that highly-methylated symmetric sites in *P. patens* are deaminated more rapidly than lowly-methylated CHH sites. In support of this conclusion, we found that only ~20% of transition mutations in *Arabidopsis* TEs^37^ were of lowly methylated CHH sites (3.3 times less than expected), the rest were of highly methylated CGs or CHGs (Supplementary Fig. 3d). Conclusively, our findings unravel the importance of the broad (high frequency) and moderate (inefficient) CHH methylation mechanism for silencing an early and transient phase in the life of TEs (Fig. 7c). Based on these results, we suggest that a mixture of CG and non-CG methylation contexts, as well as efficient and moderate methylation mechanisms are optimal combinations for genome protection and regulation under and during diverse genetic circumstances.

Regulation of gene expression by DNA methylation is the consequence of direct and indirect effects. Thus, our conclusion that non-CG methylation has a greater effect on gene regulation than that of CG methylation (Fig. 6) could be specific to *P. patens*. In contrast to genes, TEs are mainly autoregulated by DNA methylation, allowing us to make broader conclusions based on results in the first system to be analyzed for TE regulation that has comparable CG and CHG methylation patterns, that is *P. patens*. Firstly, our finding that non-CG methylation is more potent than CG methylation in silencing TEs acts as a proof-of-concept that CG methylation, despite being highly efficient and robust in many eukaryotic genomes, did not necessarily evolve as an ultimate mechanism for transcriptional silencing. This observation correlates with the involvement of CG methylation also within active chromatin regions, such as gene bodies of actively transcribed genes^1,3,38–40^. Accordingly, it is possible that non-CG methylation evolved in plants to supplement CG methylation in TE silencing and by that further allowing CG methylation to be associated with alternative genomic activities. Secondly, current knowledge suggest that all land plant species utilize CG and non-CG methylation to silence TEs, thus our finding that non-CG methylation is superior to CG methylation in TE regulation in *P. patens* could be of high biological relevance to numerous plant species, including among flowering plants, which also have vigorous non-CG methylation system, such as marihuana (*C. sativa)*, cotton (*G. raimondii*), and clementine (*C. clementina*) to name a few^6^.

## Methods

### Generation of transgenic mutant lines and growth conditions

A *ΔPpmetΔPpdnmt3b* double deletion mutant was generated by replacing the genomic region coding for PpDNMT3b in *ΔPpmet* single deletion mutant background, and a *ΔPpcmtΔPpdnmt3b* double mutant was generated by replacing the genomic region coding for PpCMT in the *ΔPpdnmt3b* single deletion mutant using the same constructs as were used for generating single mutants as described previously^17,26,27^. Mutants were verified by the absence of BS-seq reads in the knockout regions (Supplementary Figs. 3a-c).

Three independent experiments have been conducted for knocking out PpCMT in the background of Δ*Ppmet* or knocking out of PpMET in the background of the Δ*Ppcmt* line, but these attempts yielded no Δ*Ppmet* Δ*Ppcmt* double mutants, suggesting that such a combination is lethal. *P. patens* WT and mutant plants were propagated on BCD or BCDAT media^41^ at 25 °C under a 16-h light and 8-h dark regime^42^. Arabidopsis WT (Col-0) and *ddcc* mutant (*drm1-2* SALK_021316; *drm2-2* SALK_150863; *cmt2-3* SALK_012874; *cmt3-12* SALK_148381) seeds were sown on soil in pots, stratified at 4°C for 48h and grown under 22°C and long day (16h light/8h darkness) regime.

### *P. patens* plant size analysis

Images of petri dishes containing *P. patens* 27-days old plants were processed using ImageJ v1.52i (Fiji) to assess the area of each plant. Mann–Whitney test was performed for statistical evaluation of size variation between WT and double mutant plants, using GraphPad Prism software (La Jolla, CA, USA).

### BS-seq library preparation

Genomic DNA was extracted from 7 days old protonema using DNeasy Plant Mini Kit (Qiagen) according to manufacturer’s instructions. About 1 ug of purified genomic DNA was fragmented by sonication, end repaired, and ligated to custom synthesized methylated adapters. Adaptor-ligated libraries were subjected to sodium bisulfite treatment using the Methylation-Gold Bisulfite kit (ZYMO) as outlined in the manufacturer’s instructions. The converted libraries were then amplified by PCR, gel purified to select the 350-400 bp size range, quantified using real-time quantitative PCR with TaqMan probe and validated with Bioanalyzer (Agilent). The libraries were amplified on cBot to generate the cluster on the flowcell (TruSeq PE Cluster Kit V3–cBot–HS, Illumina) and sequenced as paired-end on the HiSeq 2000 System (TruSeq SBS KIT-HS V3, Illumina).

### BS-seq data analysis

Reads quality was evaluated using FastQC v0.11.8 (Babraham Bioinformatics). Identical duplicate reads and low-quality reads were filtered out using Prinseq-lite software v0-20-4^43^. BS-seq data processing was performed as described previously^44^. Shortly, we used custom Perl scripts to convert all the Cs in the ‘forward’ reads (and in the scaffold) to Ts, and all the Gs in the ‘reverse’ reads and scaffold to As. The converted reads were aligned to the converted scaffold using Bowtie2 v2.3.4.1^45^, allowing two mismatches and multimapping reads with up to ten hits. We then used Perl scripts to recover the original sequence information and, for each C (on either strand), count the number of times it was sequenced as a C or a T. We calculated fractional methylation within a 50 bp sliding window for each sequence context separately and regardless of context, for use in downstream analyses. The percent of DNA methylation change (such as in Fig. 1f) was calculated by dividing the difference in methylation level between two samples by the level of methylation in the sample with the higher methylation level. Two-dimensional density plots for visualizing CG/CHG symmetry level (such as in Fig. 3c) were generated using hist2d function with norm=LogNorm() parameter from matplotlib.pyplot Python package. For identifying DMRs, fractional methylation in 50-bp windows across the genome was compared between WT and each of the mutants. DMRs were called for windows with minimal coverage of 3 reads, and Fisher’s exact test p-value < 0.05. For hypo-DMRs and genes proximity analysis, CG and CHG DMRs with at least 40% methylation decrease between WT and mutant, and CHH DMRs with at least 20% methylation decrease between WT and mutant were considered.

### RNA-seq library preparation

Total RNA from P. patens plants was extracted from seven days old protonema using SV Total RNA Isolation System (Promega, USA) and DNase treatment (Thermo Fisher Scientific, USA). Total RNA from *Arabidopsis* plants was extracted from one month old rosette leaves using RNeasy plant mini kit (Qiagen) according to manufacturer’s instructions. The poly-A fraction (mRNA) was purified from 500 ng of total input RNA, followed by fragmentation and the generation of double-stranded cDNA. Then, Agencourt Ampure XP beads cleanup (Beckman Coulter), end repair, A base addition, adapter ligation and PCR amplification steps were performed. Libraries were quantified by Qubit (Thermo fisher scientific) and TapeStation (Agilent). The libraries were sequenced as single end reads on HiSeq 2500.

### RNA-seq data analysis

Reads quality was evaluated using FastQC v0.11.8 (Babraham Bioinformatics). Adaptor and quality trimming and polyA-tails removal were performed with TrimGalore v0.6.0 (Babraham Bioinformatics) with Cutadapt v1.15. The pre-processed reads were aligned to the reference genome and reads counts per gene/TE were obtained using STAR v2.7.0f^46^. Additionally, aligned reads were counted in a 50-bp sliding window genome-wide and normalized to library size (RPM), for use in downstream analysis, e.g. genome browser snapshots, and expression profiling across aligned TEs.

For differential expression we used voom from the edgeR/limma Bioconductor R packages, the differential expression model accounted for a possible batch effect. Genes with raw read count below 10 were omitted from the analysis. Genes and TEs with 2-fold change in expression level and false discovery rate (FDR) below 0.1 were considered differentially expressed.

### Meta analyses across genes and TEs

Meta-analysis of DNA methylation, CG/CHG/CHH site abundancy, RNA-seq reads and DMRs relatively to genes and TEs edges was performed by first aligning genes or TEs at either their 5’ or 3’ ends. Then, for each gene or TE, a score (mean methylation level or total number of sites) or the presence of DMR (1 or 0) was calculated in a sliding window (50 or 100 bp) upstream and downstream of point of alignment and a mean value per each window was calculated for all included elements. Elements were generally included when reached to either the end of their sequence, another annotated element, or the indicated selected length.

### Public datasets analysis

*A. thaliana*, *M. polymorpha*, and *O. sativa* RNA-seq and BS-seq raw reads (Supplementary Table 1) were download from NCBI SRA archive and processed as described above.

### Gene and TE annotations

Gene annotations for *P. patens* (v3.3), *A. thaliana* (Araport11), *M. polymorpha* (v3.1) and *O. sativa* (v7.0) were downloaded from Phytozome https://phytozome.jgi.doe.gov/. *P. patens* TE annotations was downloaded from CoGe https://genomevolution.org/coge/GenomeView.pl?gid=33928 and a separate annotation for intact LTR retrotransposons was kindly provided by Prof. Stefan Rensing. Information from *P. patens* Repeatmasked assembly (v3.3) downloaded from Phytozome was used to increase the resolution of LTR-TEs families. To assess the ages of LTR retrotransposons families based on LTRs similarity level, REannotate software^47^ was used for RepeatMasker output processing. LTR retrotransposons age was calculated assuming neutral mutation rate of 9.4*10^−9^ synonymous substitutions per synonymous site per year^48^.

*Arabidopsis* TE annotations were downloaded from TAIR https://www.arabidopsis.org/, TAIR10 genome release; for *M. polymorpha* and *O. sativa*, TEs were annotated *de-novo* using REPET software v3.0^49,50^.

## Data availability

New *P. patens* WGBS and transcriptome data and *A. thaliana* transcriptome data produced in this study were submitted to GEO under the accession number GSE142054.

## Acknowledgements

This work was supported by the European Research Council (ERC, 679551), Israeli Centers for Research Excellence Program of the Planning and Budgeting Committee, Israel Science Foundation (757/12), and Israel Science Foundation (1636/15) to A.Z, and Israel Science Foundation (767/09) to N.O.. We thank Dr. Vicki Plaks (University of California, San Francisco), Dr. Keith D. Harris (Tel Aviv University), and Prof. Avi Levy (Weizmann Institute) for editing and scientific comments.

## Author contributions

A.Z. and N.O. conceived and supervised the study, A.K., V.H.N., U.V.T., and K.G.H. generated and phenotyped the mutants, A.K., O.G., and E.K. conducted the genomic experiments, K.D., A.Z., and R.Y. analyzed genomic data. A.Z., K.D., N.O., and R.Y., wrote the manuscript. All authors discussed the results and commented on the manuscript.

