## Supplementary Fig. 1. for "Non-CG methylation is superior to CG methylation in genome regulation"

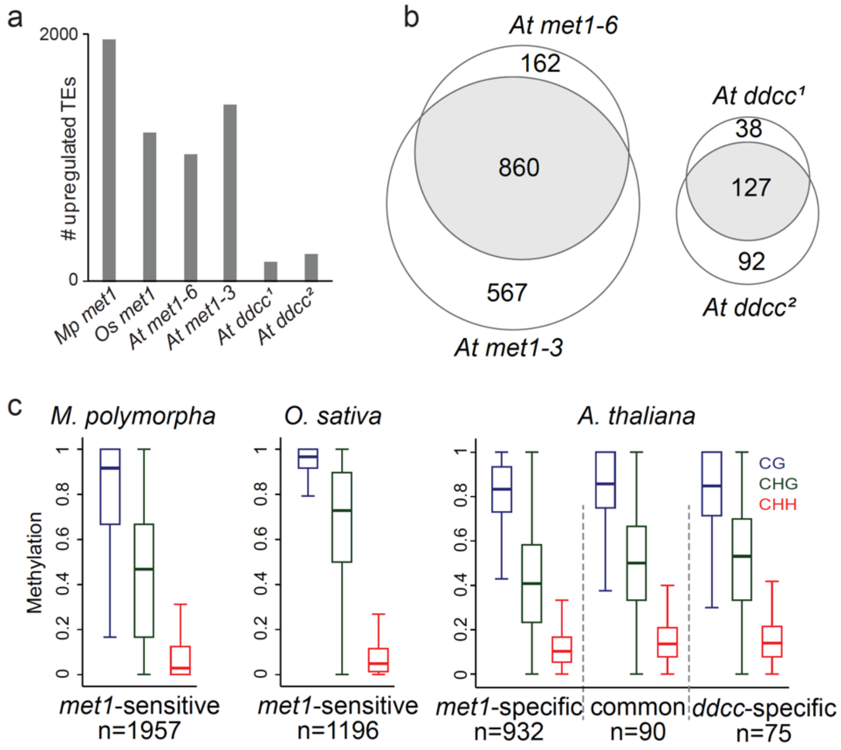


**Supplementary Fig. 1. *MET1* gene plays an important role in TE silencing in *M. polymorpha*, *O. sativa* and *A. thaliana*.**
(a) The numbers of TEs transcriptionally upregulated in the indicated *M. polymorpha*[22], *O. sativa*[23], and *A. thaliana*[32,51,52] DNA hypomethylation mutants. (b) Venn diagrams demonstrating the overlaps between upregulated TEs in two *A. thaliana met1* and two *ddcc* mutants: Left panel - the overlap between TEs upregulated in *met1-6*[32] mutant and in *met1-3*[52] mutant; right panel - the overlap between TEs upregulated in *ddcc*[51]*[1]* (by Stroud *et al*., 2014[51]) and in *ddcc*[2] (current study) mutants. (c). Box plots of CG, CHG and CHH methylation levels in WT *M. polymorpha*, *O. sativa* and *A. thaliana*, in 50 bp genomic windows overlapping TEs upregulated in the indicated mutants. *M. polymorpha* (*Mp*), *O. sativa* (*Os*), and *A. thaliana* (*At*) methylome and transcriptome public raw data (detailed in Table S1) were processed and similarly analyzed as described in Methods.
