## Supplementary Fig. 2. for "Non-CG methylation is superior to CG methylation in genome regulation"

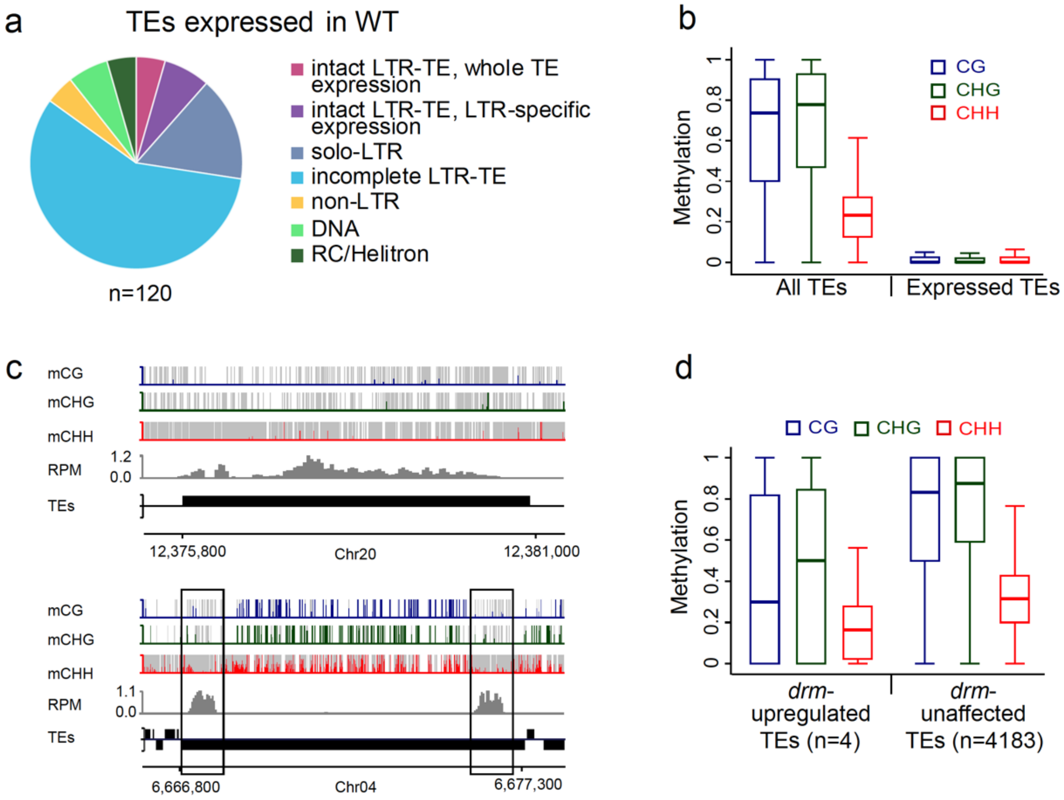


**Supplementary Fig. 2. TE expression and methylation in WT and *drm P. patens* plants.**
(a) Pie chart of expressed TEs (>0.5 RPKM in at least 2 replicates) in WT *P. patens* protonema tissue, separated by TE superfamily type or expression profile. Out of the 120 expressed TEs, over 75% are LTR retrotransposons. Among LTR superfamily, five were intact TEs with transcripts covering the entire elements. (b) DNA methylation of all *P. patens* TEs or expressed ones (as in a) in WT plants. Expressed TE sequences demonstrate low methylation levels. (c) Genome browser snapshots of unmethylated-expressed TEs. Fractional CG, CHG and CHH methylation averaged in 10 bp sliding window resolution (scales are 0-1). Grey bars represent the presence of covered sites in each methylation context in the given window. Expression is shown as normalized averaged reads coverage (RPM) in 50 bp windows. Left panel represents an intact LTR/Copia TE unmethylated and expressed throughout the entire element. Right panel, represents an intact Gypsy TE that is unmethylated and expressed exclusively within the LTR sequences (open boxes). (d) Box plots of CG, CHG and CHH methylation in 50 bp windows within the four TEs upregulated in *drm* mutant, and other TEs unaffected in *drm* but upregulated in other PpDNMT mutants.
