## Supplementary Fig. 3. for "Non-CG methylation is superior to CG methylation in genome regulation"

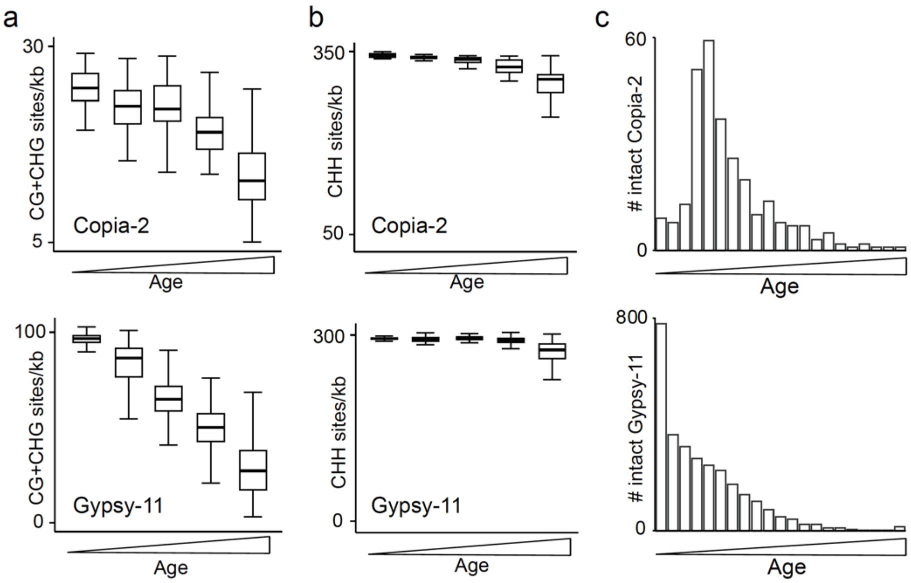


d

| Chr. | Position | Context | Methylation |
|---|---|---|---|
| 1 | 7068062 | CG | 0.985 |
| 4 | 1985681 | CG | 0.96 |
| 4 | 3688314 | CG | 0.975 |
| 1 | 7065559 | CHG | 0 |
| 2 | 6438290 | CHG | 0.745 |
| 2 | 6939643 | CHG | 0.895 |
| 3 | 15495167 | CHG | 0.785 |
| 4 | 2983511 | CHG | 0.71 |
| 5 | 6434116 | CHG | 0.845 |
| 3 | 15580586 | CHH | 0.015 |
| 4 | 4280167 | CHH | 0.24 |

**Supplementary Fig. 3. Relative more rapid depletion of symmetrically CG/CHG contexts versus asymmetric CHH in *P. patens* and *A. thaliana* TEs.**
(a) Boxplots of CG plus CHG sites frequency per kilobase of Copia-2 (upper graph) or Gypsy-11 (lower graph) over five age quantiles (ordered from young to old). TE age was calculated based on divergence between 5' and 3' LTRs of intact TEs. (b) Boxplots of CHH sites frequency per kilobase of Copia-2 (upper graph) or Gypsy-11 (lower graph) over five age quantiles. Note the relative smaller range in CHH frequency in comparison to that of CG+CHG. In Gypsy-11 CHH frequency drops only in the fourth quantile, in contrast to symmetric sites that gradually drops in each of the quantiles. (c) Histograms of intact Copia-2 (upper graph) and Gypsy-11 (lower graph) age distributions. Copia-2 and Gypsy-11 are depleted and enriched in young TEs, respectively, explaining the higher frequency of symmetrical sites in Gypsy-11 over Copia-2 (a). (d) Sequence context and methylation level of 11 cytosine transition mutations detected in TEs between 30 generations in A. thaliana[37]. The methylation level is an average of methylation of two sperm BS-seq data[53,54], i.e. germ cells in which deamination of methylated cytosines in them will be fixed in the population. The frequency of CHH sites in *Arabidopsis* TEs is 2.87 times greater than CG and CHG (combined), however among the 11 mutated cytosines only 2 were in CHH sites, the rest were mostly in highly methylated CG or CHG sites.
