## Supplementary Fig. 4. for "Non-CG methylation is superior to CG methylation in genome regulation"

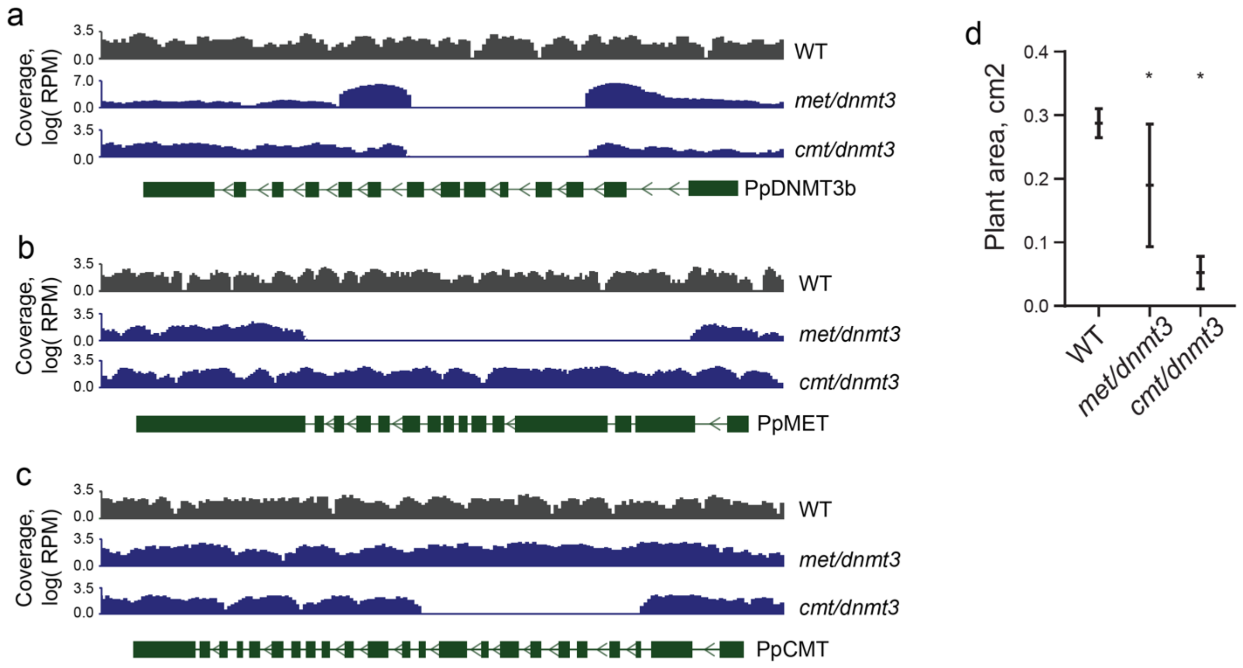


**Supplementary Fig. 4. Genotypic and phenotypic characterization of met/dnmt3 and cmt/dnmt3 mutants.**
(a-c) Genotyping of *met/dnmt3* and *cmt/dnmt3* using BS-seq data. Coverage of aligned BS-seq was quantified in 50 bp windows and normalized to library size. Genomic snapshots of PpDNMT3b/Pp3c13_8320 (a), PpMET/Pp3c11_20540 (b), and PpCMT/Pp3c6_28720 (c) shows missing coverage in the indicated mutants. (d) Scatter plot of WT, *met/dnmt3* and *cmt/dnmt3* plant sizes (n=6-13). Middle lines and whiskers show mean and SD, respectively. Asterisks indicate statistically significant size difference between mutant and WT plants (Mann-Whitney test, P-value<0.05).
