## Supplementary Fig. 5. for "Non-CG methylation is superior to CG methylation in genome regulation"

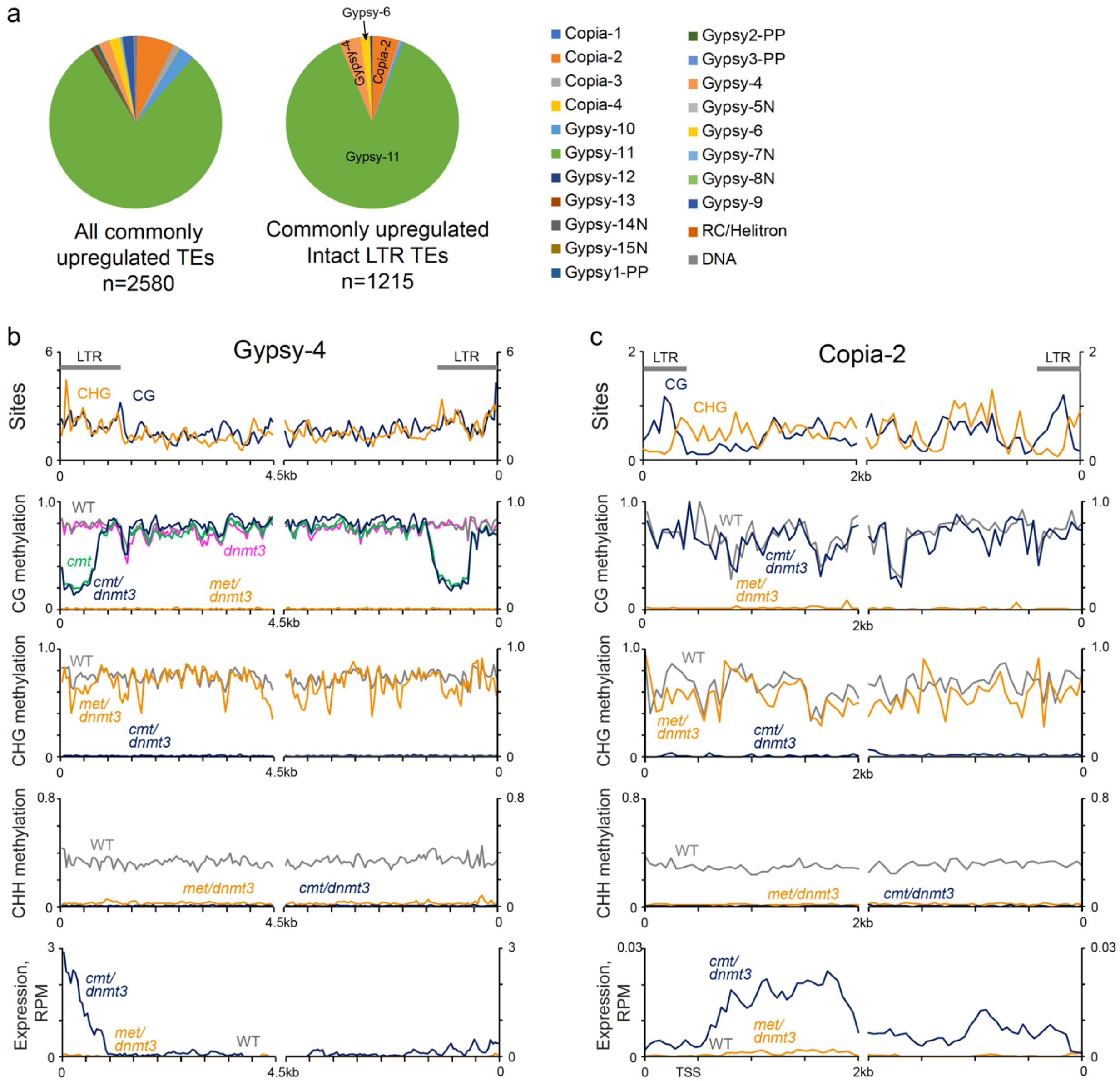


**Supplementary Fig. 5. Profiling LTR-retrotransposons commonly-upregulated in *met*/*dnmt3* and *cmt*/*dnmt3* mutants.** (a) Pie charts of all (left) and intact-only (right) commonly-upregulated TEs in *met*/*dnmt3* and *cmt*/*dnmt3* mutants, separated to TE superfamilies. The main four superfamilies are indicated on the charts. (b-c) Patterns of CG and CHG site frequency (first panel), CG, CHG, and CHH methylation in WT and indicated mutants (second to fourth panel, respectively), and expression (fifth panel) across intact commonly-upregulated Gypsy-4 (b) and Copia-2 (b) in the double mutants. Intact TEs were aligned at the 5' or the 3' ends (zero in the x-axis) and averaged for sites, methylation, and expression in a 50 bp windows. LTR sequences are marked on the top panel.
