## Supplementary Table 1 for "Non-CG methylation is superior to CG methylation in genome regulation"

| Species | Genotypes | RNA-seq | WGBS |
|---|---|---|---|
| *M. polymorpha* | WT, *met1* | PRJDB7342 | PRJDB7342 |
| *A. thaliana* | WT, *met1-6* | PRJNA176484 | PRJNA176484 |
| | WT, *met1-3* | PRJNA379224 | |
| | WT, *ddcc* | PRJNA222364 | PRJNA222364 |
| *O. sativa* | WT, *met1* | PRJNA253163 | PRJNA253162 |

**Supplementary Table 1. Summary of DNA methylation in various plants with methylome data of DNMT mutants.** NCBI bio-project accession numbers of publicly available RNA-seq and whole genome bisulfite sequencing (WGBS) data re-analyzed in this study as described in Methods section.
